# Cusp-artifacts in high order Superresolution optical fluctuation imaging

**DOI:** 10.1101/545574

**Authors:** Xiyu Yi, Shimon Weiss

**Affiliations:** Department of Chemistry and Biochemistry, University of California, Los Angeles, 90095, USA; Department of Physiology, University of California, Los Angeles, 90095, USA; California NanoSystems Institute, University of California, Los Angeles, 90095, USA; Department of Physics, Institute for Nanotechnology and Advanced Materials, Bar-Ilan University, Ramat-Gan, 52900, Israel; Lawrence Livermore National Laboratory, 7000 East Avenue, Livermore, CA 94550, USA

## Abstract

Superresolution optical fluctuation imaging (SOFI) is a simple and affordable super-resolution imaging technique, and attracted a growing community over the past decade. However, the theoretical resolution enhancement of high order SOFI is still not fulfilled. In this study, we identify “cusp artifacts” in high order SOFI images, and show that the high-order cumulants, odd-order moments and balanced-cumulants (bSOFI) are highly vulnerable to cusp artifacts. Our study provides guidelines for developing and screening for fluorescence probes, and improving data acquisition for SOFI. The new insight is important to inspire positive utilization of the cusp artifacts.

## 1. Introduction

Superresolution (SR) imaging techniques such as stimulated emission depletion microscopy (STED) [1], photo activated localization microscopy (PALM) [2], structured illumination microscopy (SIM) [3], stochastic optical reconstruction microscopy (STORM) [4] and their many derivatives have gained prominence in recent years [5–8] by providing imaging below the diffraction limit of light. Superresolution optical fluctuation imaging (SOFI) [9] is an affordable alternative to these methods. In SOFI, consecutive frames are acquired to form a movie of the imaging sample, which is labeled with stochastically blinking probes. The auto- and cross-correlations of the time trajectories of the pixel intensities are then calculated and subsequently used to construct the different-order cumulants in order to obtain high-order SOFI images. Since SOFI does not require any special hardware and is based on a simple mathematical algorithm, it has the potential to “democratize” SR imaging. The only requirement for SOFI is that the fluorescence probes used should exhibit stochastic blinking at a rate that can be captured by a camera. Quantum dots (QDs) [9], organic fluorophores (dyes) [10], fluorescence proteins [11, 12], carbon nanodots [13], and Raman probes coupled to plasmonic nanoparticles [14] have all been used for SOFI. Other forms of optical fluctuations have also been exploited for SR imaging using SOFI, such as those related to diffusion-assisted Forster resonance energy transfer [15], protein-protein interactions [16], and the diffusion of nonblinking probes [17]. The large variety of probes available and the various implementations of SOFI suggest that it may be useful in a variety of applications.

The resolution enhancement of SOFI is manifested by the reduced width of the point spread function (PSF) in the reconstructed SOFI image. Theoretically, the PSF width for a *n*^*th*^-order SOFI image is reduced by a factor of *n*^*1/2*^ as compared to that of the PSF in the original acquisition. When deconvolution or Fourier reweighting [18] is combined with the estimation of the system’s optical PSF [18], an additional enhancement of *n*^*1/2*^ can be realized, bringing the total theoretical resolution enhancement factor to *n*. Moreover, in principle, the resolution could be improved even further by increasing the SOFI order, *n*.

However, in practice, the resolution enhancement of high-order SOFI is limited by two fundamental issues. The first issue is the nonlinear expansion of the dynamic range of the pixel intensities in high-order (order > 2) SOFI images. This increase in the dynamic range renders the fine details in SOFI images imperceptible. The balanced SOFI (bSOFI) [19] method has been developed and adopted widely [20] for solving the first problem. This method involves calculating the balanced cumulants, which compensate for the expanded dynamic range of the pixel intensities in high-order SOFI images. However, the application of bSOFI in the case of high-order cumulants (> 4) has rarely been reported and generally results in artifacts in practice. The second issue, which was mostly ignored in the past but was identified in this study, is attributable to the positive and negative oscillations of the cumulants (Fig. 1). More specifically, the boundaries between the negative and positive regions result in artifacts (we have labeled these as “cusp artifacts”). We found that cusp artifacts are intrinsic to high-order SOFI cumulants due to finite and inhomogeneous blinking profiles with limited signal to noise ratio. Subsequent post-processing steps that rely on the high-order SOFI cumulants would probably be adversely affected by these artifacts, such as deconvolution algorithms with positivity constraints (e.g., MATLAB’s “deconvlucy” and “deconvblind” functions), and high-order bSOFI reconstructions, which assumes perfect deconvolution.

**Fig. 1.**
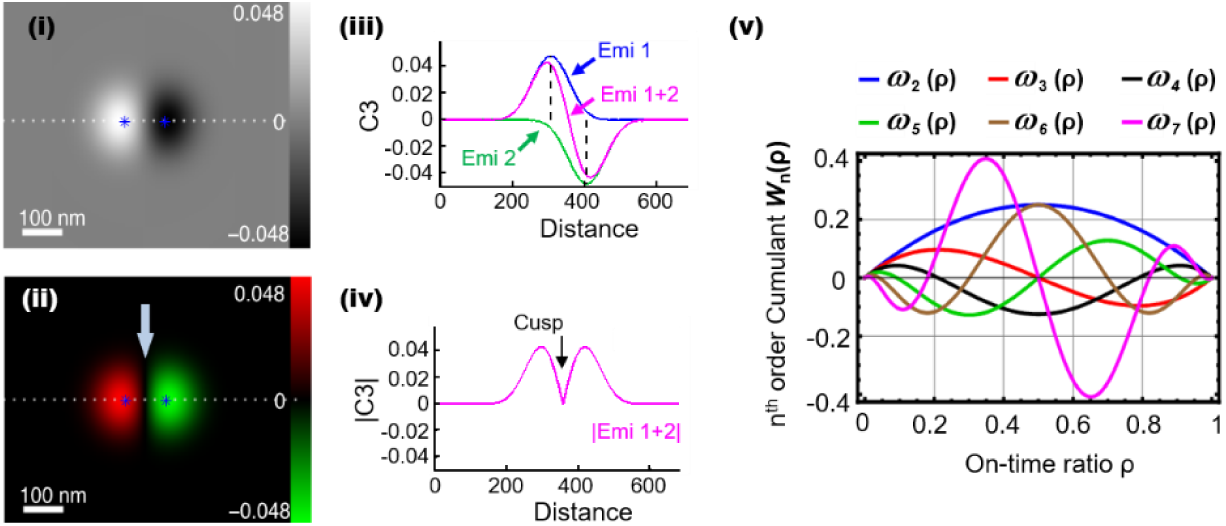
Demonstration of cusp artifact. Panels (i) and (ii) show two representations of same theoretical 3^rd^-order SOFI cumulant image of two emitters. (i) shows grayscale display, while (ii) shows g/r color code, with red (left lobe) representing positive cumulant values and green (right lobe) representing negative cumulant values; both have dynamic range of −0.048 to 0.048. On-time ratio of left emitter was set to 0.4 and that of right emitter to 0.6. Panels (iii) and (iv) show cross-sectional plots at dashed lines in (i) and (ii), respectively. Panel (v) shows plots of cumulants *ω*_n_ as function of ρ, with n = 2–7. All cumulants (n > 2) oscillate between negative and positive values, and the number of zero crossings increases with the increase of cumulant order. The total number of zero crossings is always (n-2). As can be seen, *ω*_3_(0.4) > 0 and *ω*_3_(0.6) < 0, corresponding to virtual brightnesses shown in (i) and (ii). Cusp artifact for this example is highlighted by arrow in (iv).

We identified cusp artifacts in experimental data, and analyzed them both theoretically and through simulations of various conditions. Our study shows that the cusp artifacts are originated from the nature of a mixture of positive and negative “virtual emitters” (explained in section 3) in the high order cumulant image, and the insights gained regarding such mixed negative and positive values can serve as guidelines for avoiding the cusp artifacts, and inspire new applications.

Concerning avoiding the cusp artifacts, our study suggests emitters with uniform photophysical parameters (blinking and bleaching) should be used; long movies should be acquired to ensure sufficient statistics in the acquisition, and the blinking statistics of the fluorophores should be tuned to be free of cusp-artifacts (see Fig. 1). Besides, bleaching should be minimized and bleaching correction [14] should be performed on the data set. Cusp artifacts are more pronounced for high-order cumulant reconstructions, where the expansion of the dynamic range degrades the quality of the reconstructed image. We used a cusp-artifact-independent method to compress the dynamic range of the pixel intensities (labeled as *ldrc*; the details are provided in [21]). We compared the obtained results with those of bSOFI reconstructions. The analyses of the theory, simulations and the experimental data all showed that the bSOFI algorithm fails to faithfully reconstruct the true image at high orders when influenced by cusp artifacts. We also examined and compared a moments-reconstruction approach to cumulants-reconstruction approach. While more details on moments reconstruction are provided in [21], we show here that even-order moments are immune to cusp artifacts and tend to smoothen the features-of-interest, while still allowing for some degree of resolution enhancement (over the diffraction limit, with the limit being 2^1/2^-fold for the *n*^th^-order moment and an additional *n*^1/2^-fold when deconvolution is performed, resulting in an overall theoretical resolution enhancement of (2*n*)^1/2^). Therefore, the proposed method has significant potential for use as a practical (but mathematically nonrigorous) solution for avoiding cusp artifacts. Compared to the case for bSOFI, the even-order moments combined with *ldrc* yield significantly more faithful images up to the 6^th^ order. (Details are presented in the accompanying manuscript [21]). We showed both theoretically and experimentally that cusp artifacts could be avoided by using even-order-moment reconstruction [21]. With the provided insights about the nature of mixed positive and negative values in the cumulant image, new methods could be developed to decipher the underlying physics through the statistics revealed by the nature of the cumulants by combining multiple orders of cumulants and solve the underlying blinking statistics altogether as a global inverse problem.

The rest of the manuscript is organized as follows: in Section 2, we briefly review the underlying theory of SOFI. In Section 3, we introduce the mathematical concept of “virtual emitters” and “virtual PSF” for high-order SOFI images (to be used in the subsequent sections). In Section 4, we present a theoretical explanation of cusp artifacts. In Section 5, we examine the conditions that result in cusp artifacts. Next, in Section 6, we evaluate the adverse effects of cusp artifacts on balanced cumulants and post-processing deconvolution algorithms. Further, we show that cusp artifacts can be eliminated completely by utilizing even-order moments (instead of cumulants) for image reconstruction. In Section 7, we compare the performances of the various algorithms using real data. Finally, we conclude the manuscript by discussing the implications of our findings in Sections 8.

## 2. Review of SOFI theory

A brief review of SOFI theory is given below. For SOFI reconstruction, a stack of frames (a movie) is obtained using a simple wide-field imaging system. The sample is labeled with stochastically blinking probes. Each point emitter (probe) in the sample plane is imaged onto the camera plane via the optical imaging system. Further, owing to the diffraction limit of light, the intensity distribution of imaging system takes the shape of the PSF. The signal captured at a given camera pixel located at 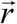 can be expressed as follows (excluding the binning effects due to pixilation):

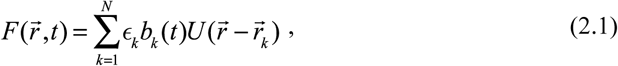

where 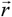 is the location of the pixel in the imaging plane, *N* is the total number of emitters, *k* is the emitter index, 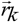 is the location of the *k*^th^ emitter, є_k_ is the brightness of the *k*^th^ emitter when it is in the “on” state, and *b*_*k*_(*t*) is the blinking time trajectory of the *k*^th^ emitter. *b*_k_(*t*) = 1 when the emitter is in the “on” state, and *b*_*k*_(*t*) = 0 when the emitter is in the “off” state. 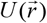 is the PSF of the imaging system, which is determined by the optical setup as well as the emission wavelength of the emitters.

In SOFI, the temporal average of each pixel’s time trajectory is subtracted from the signal, such that only the fluctuations (around zero) are considered:

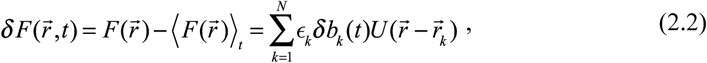

where < >_t_ represents the time-average operator, and δ*b*_k_(*t*) represents the fluctuations in the blinking time trajectory *b*_*k*_(*t*). The cumulants of 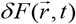 can then be calculated. In the case of a 2^nd^-order cumulant, the cumulant (C_2_) is equivalent to the correlation function:

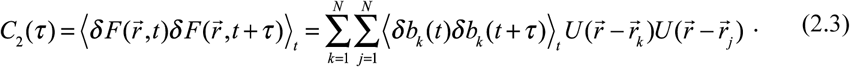

Assuming that the emitters blink independently, the temporal cross-correlation between the blinking trajectories of two emitters is zero. Only the auto-correlation of a single emitter trajectory with itself yields nonzero values:

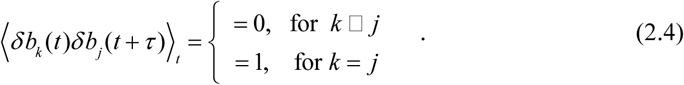

So, Eq. (2.3) becomes:

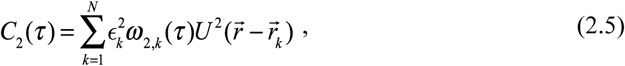

where *ω*_2,*k*_(*τ*) is the 2^nd^-order cumulant of δ*b*_*k*_(*t*). The derivation above can be extended to higher-order cumulants as well [22]. Notice that the cumulants are additive [22]. Hence, the n^th^-order cumulant of 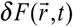 can be expressed as the sum of the cumulants of the individual emitters:

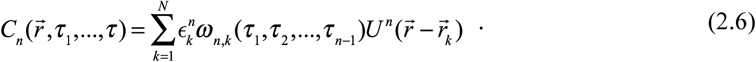

Where *ω*_*n,k*_ (*τ*_1_, *τ*_2_, …, *τ*_*n*_) is the n^th^-order cumulant of δ*b*_*k*_(*t*) and can be simplified as *C*_*n*_[*δb*_*k*_(*t*)] when the time lags are not specified.

## 3. The “virtual-emitter” interpretation for high-order SOFI

In order to better facilitate the analysis and discussions about cusp-artifacts, we propose a conceptual metaphor to interpret the physical meaning of high order SOFI cumulants based on its mathematical description (Eq. (2.6)). We note that Eq. (2.1) (which describes the direct observation image) and Eq. (2.6) (which describes the n^th^-order SOFI cumulant) share a common form. The original image (Eq. (2.1)) is formed by the summation of weighted signal from all the emitters, where the weighting factor 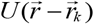 describes the weight determined by the PSF 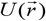 and the distance between the pixel location 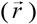 and the emitter location 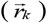 in the sample plane, and the term *ϵ*_*k*_*b*_*k*_(*t*) describes the time dependent brightness profile of the *k*^th^ emitter. Eq. (2.6) is of the same form as Eq. (2.1) with two terms replaced: The point spread function term 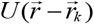 in Eq. (2.1) is replaced by 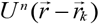 in Eq. (2.6), and the time dependent brightness term δ*b*_*k*_(*t*) in Eq. (2.1) is replaced by 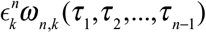 in Eq. (2.6). Therefore, Eq. (2.6) can be perceived as describing an image acquired by an optical system with a PSF *U*^n^ with signals generated by sample with ‘virtual’ emitters located at the same locations of the real emitters, but with emitter brightness described by 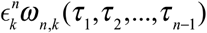. We address the term 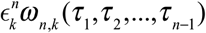 in Eq. (2.6) (which corresponds to the brightness term in Eq. (2.1)) as the “virtual brightness”, and the term 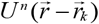 (which corresponds to the PSF term in Eq. (2.1)) as the ‘virtual PSF’. Notice that the emitter locations 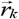 are not changed, i.e. virtual emitters share the same locations as the real emitters in the sample. Additionally, the virtual PSF is the original PSF raised to the power of *n*. In cases when the PSF can be estimated by a three-dimensional Gaussian function, the width of the PSF is reduced by a factor of *n*^1/2^, as shown below:

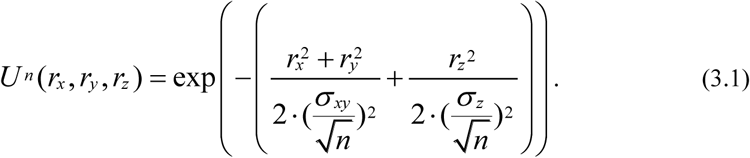

where σ_xy_ and σ_z_ are the respective widths of the original PSF in the xy- and z-planes. The reduction of the virtual PSF widths provides the basis for resolution enhancement in the SOFI cumulants (in all three dimensions). It is worth noticing that the virtual PSF is the n^th^ power of the real PSF regardless of the profile of the real PSF.

In summary, a SOFI cumulant image is equivalent to an image captured with a virtual microscope that has a virtual PSF being the nth power of the original PSF (i.e., reduced width and increased resolution with Gaussian PSF estimation), and the captured signal is created by virtual emitters that are located at exactly the same locations as the real emitters, but with altered brightnesses (i.e. virtual brightnesses) that are the cumulant of the blinking profile of each emitter (and can exhibit either positive or negative values).

We will demonstrate in the following sections that *ω*_*n,k*_(*τ*_1_, *τ*_2_, …,*τ*_*n*−1_) exhibits negative and positive values due to two main causes: blinking behavior and finite acquisition time.

## 4. Origin of cusp artifacts

For simplicity, we discuss here a noise free condition without bleaching, and assume a two-state blinking profile *b*(*t*) for the real emitters, with *b*(*t*) = 1 indicating the on state and *b*(*t*) = 0 indicating the off state. Further, we define ρ_*k*_ as the fraction of time the emitter spends in the on state during the signal acquisition step (ρ_*k*_ is the on-time ratio). Adjacent emitters could have different ρ_*k*_ values (in principle) owing to the following reasons: (i) finite acquisition time (i.e., not reaching statistical significance to represent the underlying statistical process); (ii) subtle changes in the local microenvironments; and (iii) inhomogeneity in the photophysical properties of the emitters (for example, quantum dots). And the cumulants of *b*_k_(*t*) can be expressed as a function of ρ_*k*_. If the acquisition time is long enough, and if the blinking behavior can be statistically described by a Poisson process (or Bernoulli statistics), ρ_*k*_ converges to

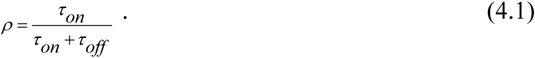

In cases when Eq. (4.1) is not applicable, such as with pure power-law blinking statistics when ρ_k_ does not converge [23], or with insufficient acquisition time to ensure statistical significance, or when the blinking behavior follows a different model, different formulas of on-time ratio ρ_k_ can be deduced. Regardless, 0≤ρ_k_≤1 holds for any two-state blinking model. Therefore, for generality, we present the following discussions in the form of functions of ρ_k_. If, for simplicity, we set all the time lags equal to zero in the cumulant calculations, we find that the cumulants with orders higher than two will exhibit positive-negative oscillations as a function of ρ_k_. The expressions for the first six cumulants (2^nd^ order to 7^th^ order) as functions of ρ_k_ are given as follows (the derivations are given in Appendix 1 [28]):

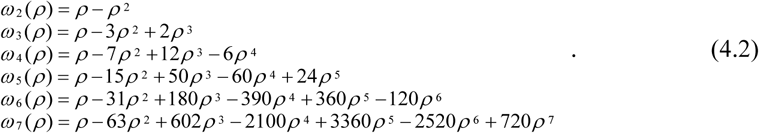

Here, the emitter index, *k*, and the time lags (τ_1_, …, τ_n−1_) have been eliminated to simplify the notation.

Fig. 1v shows cumulants of different orders as functions of the on-time ratio, ρ. The higher-order cumulants (> 2^nd^ order) clearly exhibit positive-negative oscillations as a function of ρ. If the virtual brightnesses of two nearby virtual emitters have opposite signs (Fig. 1i and Fig. 1ii), the corresponding amplitude cross-section of the SOFI amplitude image (consisting of the convolution of the virtual emitters with the virtual PSF) will exhibit positive and negative lobes (Fig. 1iii). The absolute value of this cross-section (determining this is a common step in displaying an image) exhibits a cusp (Fig. 1iv, arrow). We have therefore labeled such artifacts as “cusp artifacts.”

In order to better demonstrate cusp artifacts, we performed simulations for three adjacent blinking emitters with the same on-state brightness, but with on-time ratios of 0.831, 0.416, and 0.103 respectively (Fig. 2). The simulations show clearly that high-order cumulants can take negative values that coexist with the positive values, leading to cusp artifacts.

**Fig. 2.**
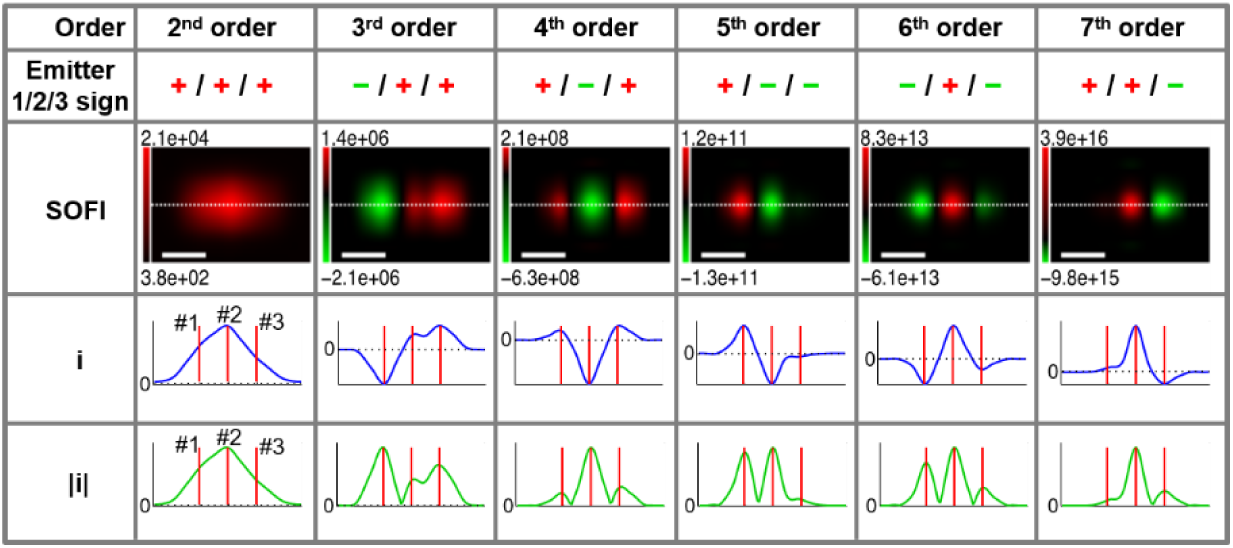
Demonstration (through simulations) of cusp artifacts for three adjacent blinking emitters spaced equally along line (spacing of 193 nm between nearest neighbors). Simulation parameters were: emission wavelength = 800 nm; numerical aperture (NA) = 1.4; and pixel size = 93.33 nm. For blinking statistics (simulated by Monte Carlo method), we set ρ to 0.831, 0.416, and 0.103 for emitters 1, 2, and 3 respectively. Orders of cumulants are shown in top row. Signs of virtual brightness values, as predicted in Fig. 1v, are denoted by (+/-) signs (second row). Third row shows simulated SOFI images for cumulants with orders 2–7 (left to right). Fourth row (i) shows cross-sections for dotted white line in second row (in blue; red line denotes emitter positions). Fifth row shows absolute value cross-sections of |i| for dotted white line in second row (in green; red line denotes emitter positions). Sign oscillations and cusp artifacts are evident, except for 2^nd^ order. Scale bars: 280 nm.

Moreover, when displaying gamma-corrected [24] experimental high-order SOFI-processed images (before performing deconvolution or Fourier reweighting), cusp artifacts can be observed readily for orders greater than two, as can be seen from the gray/red boundaries in Fig. 3.

**Fig. 3.**
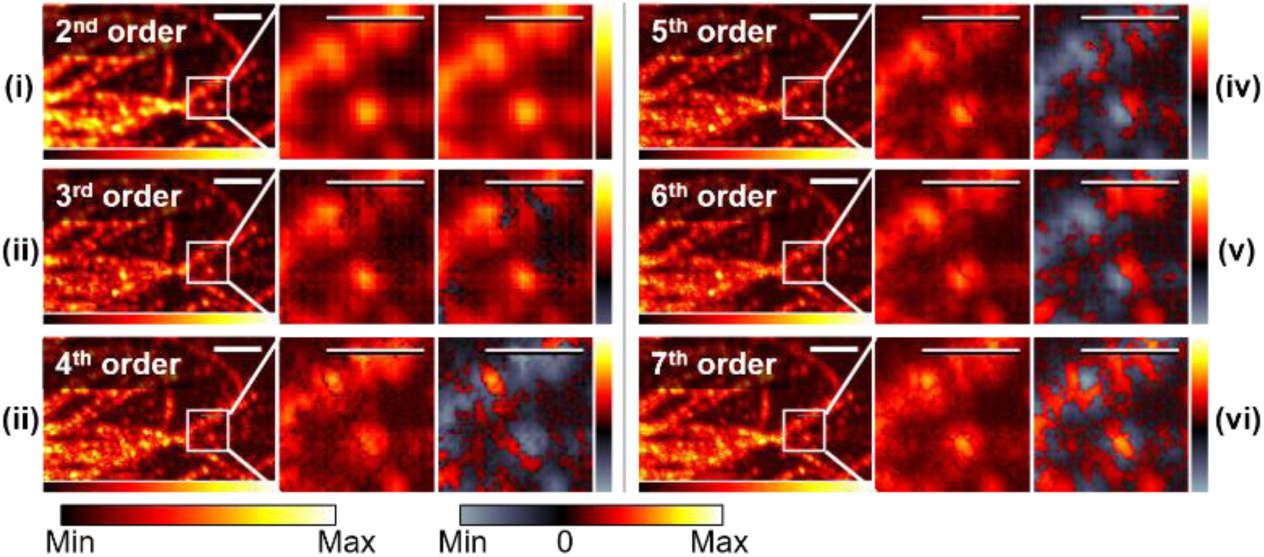
Gamma-corrected high-order SOFI-processed experimental images displaying cusp artifacts. Fixed HeLa cells were labeled with QDs (emission wavelength = 800 nm) by immunostaining using primary antibody (eBioscience, Cat#: 14-4502-80) and secondary antibody conjugated to QD800 (ThermoFisher Scientific. Ref#: Q11071MP). Total of 2000 frames (exposure time of 30 ms) were processed to obtain SOFI cumulants with up to 7^th^ order using both auto- and cross-correlations. In order to better illustrate source of cusp artifacts, final SOFI processing steps of deconvolution and Fourier reweighting were skipped. Each SOFI image of particular order is presented in three panels: large field-of-view (left), magnified absolute value SOFI image of box area (middle), and magnified positive/negative values SOFI image of box area (right). Positive/negative domains are color coded separately as shown by color bars for each panel, with color scheme shown at bottom. Cusp artifacts can be seen clearly for cumulants of orders greater than two: spatial distributions of cusps for cumulants of different orders differ and are located at boundaries between positive and negative domains. Scale bars: 3.2 μm (left) and 1.6 μm (middle/right). Image intensities are displayed with gamma correction to highlight the cusps, therefore resolution enhancement is not evident. Gamma values are the multiplicative inverse of the cumulant order. More comprehensive displays are available in Appendix 5 [28].

## 5. Exploration of parameter space of cusp artifacts

In order to better understand the prevalence and significance of cusp artifacts in SOFI reconstructions, we performed a series of realistic simulations to explore the relevant parameter space. A simulator was developed that propagates the emissions of point emitters in the sample plane onto the detector (camera) plane using the Gibson-Lanni PSF model [25]. The point emitters’ time blinking trajectories were simulated to blink according to Poisson statistics. Poisson noise was simulated as described in the accompanying manuscript [21]. Actual experimental background noise was recorded using an electron-multiplying charge-coupled device (EMCCD) camera and added to the simulated movie. Next, the SOFI cumulants of up to the 7^th^ order were calculated and analyzed. Long (20,000 frames) simulations were performed to ensure that the blinking trajectories exhibited suitably high statistical significance. Details of the simulator are given in the accompanying manuscript [21] and posted on a public GitHub account as SR_Simu3D [26].

In the first set of simulations (“Simulation-1”), we simulated four different populations of emitters (P1, P2, P3, and P4) with different ρ distributions (Fig. 4i, dashed red, blue, black, and green curves), whose ranges were as follows: 0.49 ≤ ρ ≤ 0.51 for P1, 0.53 ≤ ρ ≤ 0.87 for P2, 0.11 ≤ ρ ≤ 0.93 for P3, and 0.05 ≤ ρ ≤ 0.95 for P4 (Fig. 4i). Comparing these distributions to Fig. 1v allowed us to predict the signs of the resultant virtual brightnesses (Fig. 4ii). A simulated filamentous morphology was then populated with emitters from either P1, P2, P3, or P4. The resulting simulated movies were SOFI-processed up to the 7^th^ order. The signs of the virtual brightnesses of the virtual emitters (defined in Section 3) were mostly in keeping with the predictions in Fig. 4ii for the different orders, with the exception of the out-of-focus P2 emitter regions (details given in Fig. S1) and the 4^th^-order cumulant for P3. As can be seen from Fig. 4i, for P1, ρ is distributed in a region with positive lobes for the 2^nd^- and 6^th^-order cumulants, a negative lobe for the 4^th^-order cumulant, and positive/negative transition regions for the 3^rd^-, 5^th^ -, and 7^th^-order cumulants. This means that all P1 virtual emitters will exhibit positive virtual brightnesses for the 2^nd^- and 6^th^-order cumulants, negative virtual brightnesses for the 4^th^-order cumulant, and both negative and positive virtual brightnesses for the 3^rd^-, 5^th^-, and 7^th^-order cumulants. We could therefore predict that P1 will exhibit cusp artifacts for the 3^rd^-, 5^th^-, and 7^th^-order cumulants. Small discrepancies could be introduced by the Poisson noise in the signal as shown by the small green region for 6^th^-order cumulant image for P1. Similarly, for P2, ρ is distributed in a region with positive lobes for the 2^nd^- and 5^th^-order cumulants, negative lobes for the 3^rd^-, 4^th^-, and 7^th^-order cumulants, and positive/negative transition regions for the 6^th^-order cumulant. We could therefore predict cusp artifacts for the 6^th^-order cumulant of P2. Discrepancies can be introduced by Poisson noise in the signal (5^th^-order image for P2). Additionally, for P3 and P4, ρ is broadly distributed in the positive/negative transition regions for all the cumulants with orders higher than two and is purely positive only for the 2^nd^-order cumulant. Therefore, we predicted that P3 and P4 would exhibit cusp artifacts for all the cumulants with orders higher than 2. However, as shown in Fig. 4iii, the 4^th^-order of P3 exhibits pure negative virtual brightnesses, which can be attributed to the fact that the percentile population with positive virtual brightnesses in P3 for the 4^th^-order cumulant was too small. Therefore, the signal is canceled by the large population with negative brightnesses. This is confirmed in the case of P4, where we increased the population with positive virtual brightnesses for the 4^th^-order cumulant, and the cusp artifacts is as predicted.

**Fig. 4.**
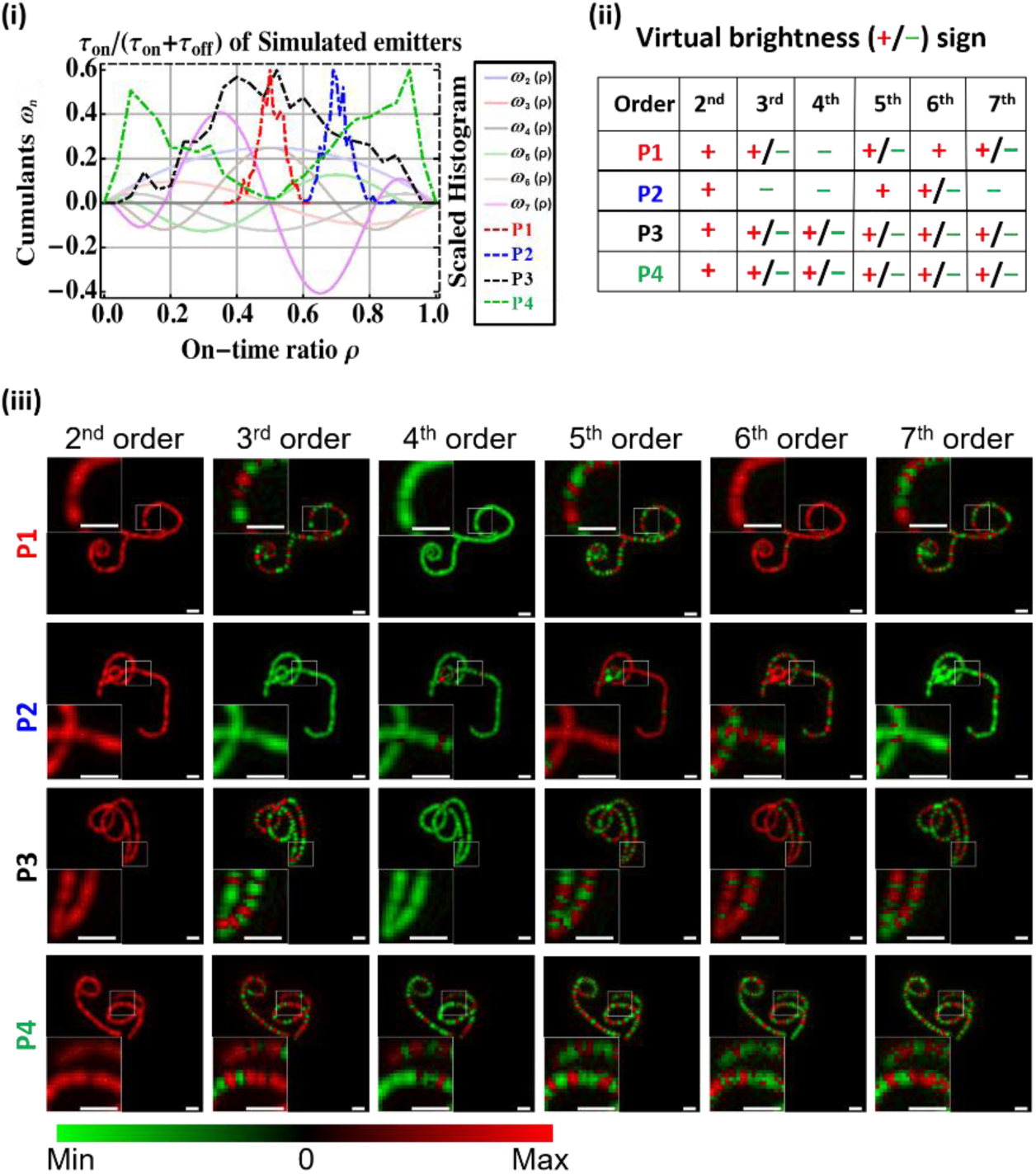
Simulations (‘Simulation-1’) showing dependence of cusp artifacts on blinking statistics. (i) Four different populations of simulated emitters with different distributions of τ_on_ and τ_off_ values, yielding four different distributions of ρ values (P1, P2, P3, and P4; dashed red, blue, black, and green curves respectively) are plotted on top in Fig. 1v. (ii). Predicted signs of calculated virtual brightnesses for P1, P2, P3, and P4 for cumulants with orders of 2 to 7. (iii) SOFI processing of simulated data. Simulated filamentous morphology was populated with emitters from either P1, P2, P3, or P4. Signs of virtual brightnesses (red for positive, green for negative) of virtual emitters are mostly in keeping with predicted ones in (ii) for different orders, with exception of out-of-focus (P2) regions (details are given in Appendix 2 [28]) and case with imbalanced population of virtual brightnesses, where smaller portion is more attenuated (4^th^-order cumulant of P3 contains 14.5% positive virtual emitters, and positive virtual brightnesses are attenuated by small amplitude of cumulant). Scale bars: 933 nm. Image intensities are displayed with gamma correction, with gamma value equals to multiplicative inverse of the cumulant order.

In the simulations, we assumed that the blinking phenomenon is governed by a Poisson process, that is, the photons emitted from an individual emitter exhibit a telegraph-noise-like time trajectory. If enough data (i.e., a large number of movie frames) with sufficient statistical significance were to be collected, it would be seen that the estimator for the “on”-time ratio, ρ, would converge to *τ*_*on*_ / (*τ*_*on*_ + *τ*_*off*_) (Eq. (4.1)). As the total number of frames in the simulation is reduced, the actual (calculated) value of ρ starts to deviate from its estimated value. This can lead to unexpected cusp artifacts for cumulants of certain orders in the regions that are predicted to be free of these artifacts. Appendix 2 [28] shows that the higher the SOFI order, the higher the number of frames needed for SOFI processing to realize the theoretically predicted cusp-artifact-free images. Moreover, since the high-order virtual brightnesses exhibit greater oscillations, they are more susceptible to heterogeneities in the photophysical properties of the emitters and hence more vulnerable to cusp artifacts.

The next set of simulations (“Simulation-2”) were performed to examine the effects of photobleaching and noise on cusp artifacts (Fig. 5). The time trajectories simulated during Simulation-1 for P1 were stochastically truncated (using Poisson bleaching statistics) to simulate the bleaching events (see Appendix 3 [28] and Fig. 5i - iii). The predicted signs of the virtual brightnesses of the emitters in P1 are shown on top in the first row of Fig. 5iv. As can be seen from Fig. 5iv, when there is no bleaching correction (see below), the signs of the virtual brightnesses displayed in the SOFI images deviate from those predicted. This is because bleaching causes ρ to deviate from its estimated value (Eq. (4.1)), as the “bleached” state is equivalent to an ultralong “off” state, causing a decrease in the ρ values of the emitters and vary from the theoretical predicted ρ value shown in Eq. (4.1). The “bleached” state also affects the assumption of independence of the blinking trajectories of the different emitters, rendering the predictions less reliable. Next, we used a bleaching correction algorithm [14] with minor modifications (Appendix 3 [28]) on the simulated data by dividing the movie into individual blocks of frames, wherein each block had a signal decrease (from the beginning to the end of the block) of f_bc_ = 1% of the overall signal decrease (from the beginning to the end of the whole movie; f_bc_ is called the “bleaching correction factor”). The final cumulant reconstruction was computed as the average of the cumulants of all the time blocks of the same cumulant order. It can be seen that the bleaching-corrected reconstructions (second row in Fig. 5iv) exhibit the predicted virtual brightness signs. Next, we examined the effects of the signal-to-noise ratio (SNR) on the bleaching correction. The noise of the EMCCD camera was recorded and added to the simulated bleaching data with altered signal levels (as described in Appendix 3.2 [28]) to ensure SNR levels of 1.47 dB (third row) and −1.33 dB (bottom row). The results of the simulations with bleaching and background noise were SOFI-processed up to the 7^th^ order. As can be seen in Fig. 5, the noise severely degrades the quality of the images in most of the cases, especially for the cumulant orders with a large proportion of positive virtual emitters. However, if the emitters’ blinking parameters were to be tuned to have either purely or primarily negative virtual brightnesses (as for the 4^th^-order cumulant in the third row, which has pure negative virtual brightnesses, and the 7^th^-order cumulant in the third row, which primarily has negative virtual brightnesses), a significant enhancement in contrast would be achieved (as shown for this set of simulations). If the majority of the virtual emitters are negative, the negative signal would still create a negative contrast against the positive noise background (as can be seen for the 7^th^-order cumulant in the third and fourth rows). We would like to note here that the cumulants of the background noise were positive for all orders higher than two; this was because the higher-order (>2) cumulants of a random variable that follows a Poisson distribution remain constant (Appendix 4 [28]).

**Fig. 5.**
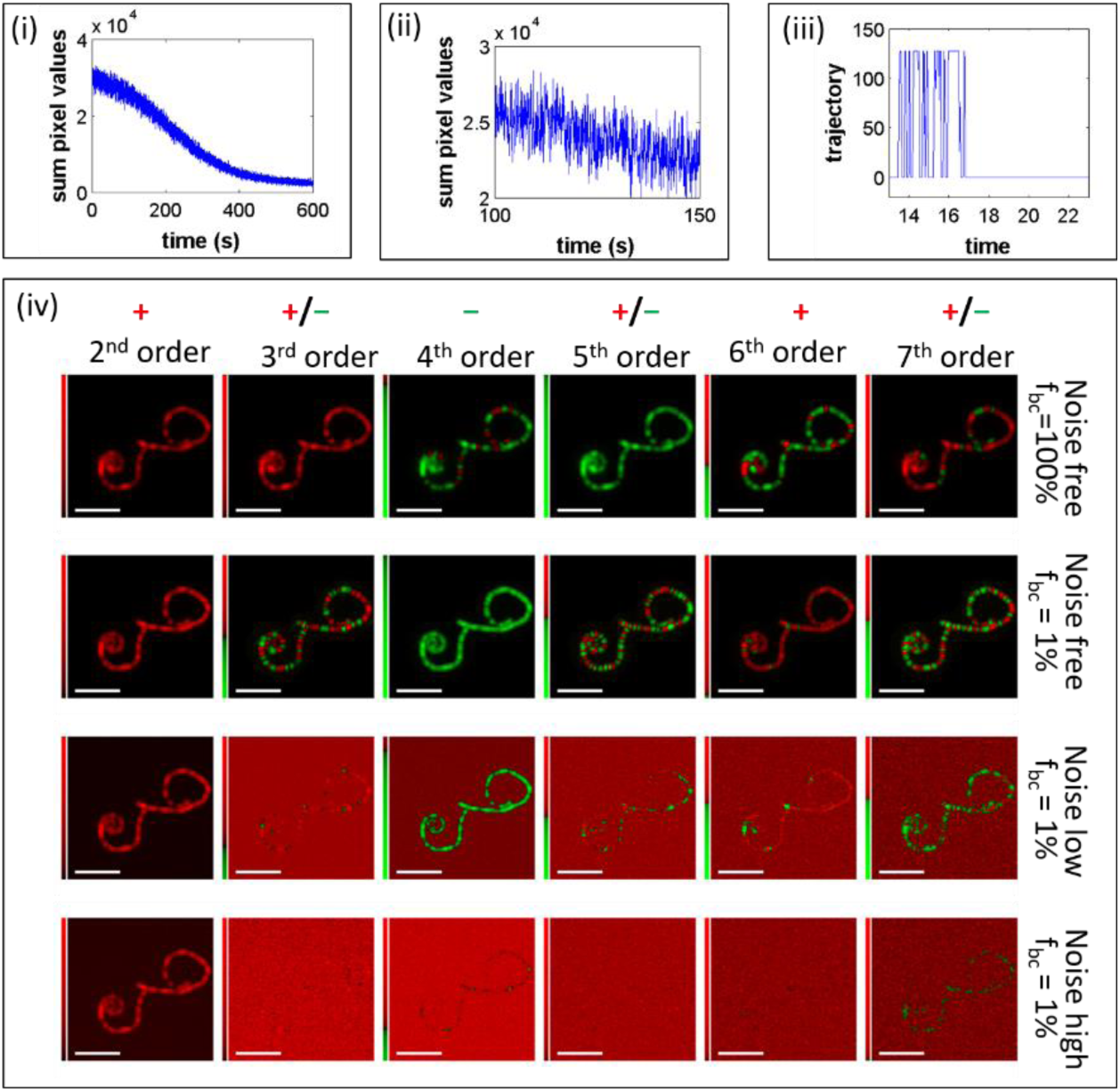
Simulations (‘Simulation-2’) to evaluate effects of bleaching and noise on cusp artifacts. (i) Total signal, summed over all pixels, as function of movie frame (time), after bleaching operator was used (before addition of background noise). (ii) Magnified version of curve in (i), showing intensity fluctuations. (iii) Example of single-emitter blinking trajectory that was “bleached” at approximately ~17 s. (iv) Array of SOFI cumulant images for orders 2^nd^ to 7^th^ as functions of noise. Top row shows simulated images with no bleaching. Signs of virtual brightnesses are in keeping with predictions in Fig. 4i (bleaching correction factor, f_bc_, of 100% means there was no bleaching correction). Second row shows simulated images for bleaching correction factor, f_bc_, of 1% (see text). Bleaching correction algorithm was effective in restoring absolute value of virtual brightness distribution but not brightness signs. Bleaching correction protocol changed signs of cumulants, resulting in rapid sign changes in cases of 3^rd^, 5^th^, and 7^th^ (odd) orders. Real background noise (recorded as empty frames with EMCCD camera) when added to simulated bleaching data severely degrades quality of images (background noise is always positive). If emitters’ blinking statistics yield pure negative virtual brightnesses, as shown for 4^th^-order cumulant in third row, significant enhancement in contrast results. Scale bars: 2.8 μm. Image intensities are displayed with gamma correction, with gamma value equals to multiplicative inverse of the cumulant order.

Here, we would like to highlight the trade-offs between the bleaching correction factor (f_bc_), noise level, and statistical significance of the blinking trajectories (details available in Appendix 3 [28]). When we decreased f_bc_, the block size decreased, and the total number of bleaching events occurring within each block decreased as well. Thus, the bleaching effect was suppressed to a greater degree. However, because the smaller f_bc_ led to a decrease in the degree of distinctiveness between the blinking events and noise within each block, the construction of the cumulants became more vulnerable to noise. In addition, a smaller f_bc_ reduces the statistical significance of the blinking trajectories within each block and increases the total number of blocks for constructing the final cumulant. Thus, these two factors counterbalance each other in terms of the effects of f_bc_ on the overall statistical significance of the blinking trajectories.

In the simulations discussed above (Simulation-1 and Simulation-2), the spatial distribution of the ρ value of the different emitters was random. In Fig. 6, we show the results of another set of simulations (“Simulation-3”) in which the ρ values were varied slowly over the range of 0.01 to 0.99 in increments of 0.01 across the spatial field of view, and the SOFI cumulants of the 2^nd^ to 7^th^ orders were calculated. As can be seen from the figure, the number of zero-crossing nodes increased with the order of the cumulants, with the positive and negative domains being in good agreement with the ground-truth predictions. Further, the spatial variations in the blinking statistics can be seen visually as green/red segments (with the cusps concentrated at the segments boundaries).

**Fig. 6.**
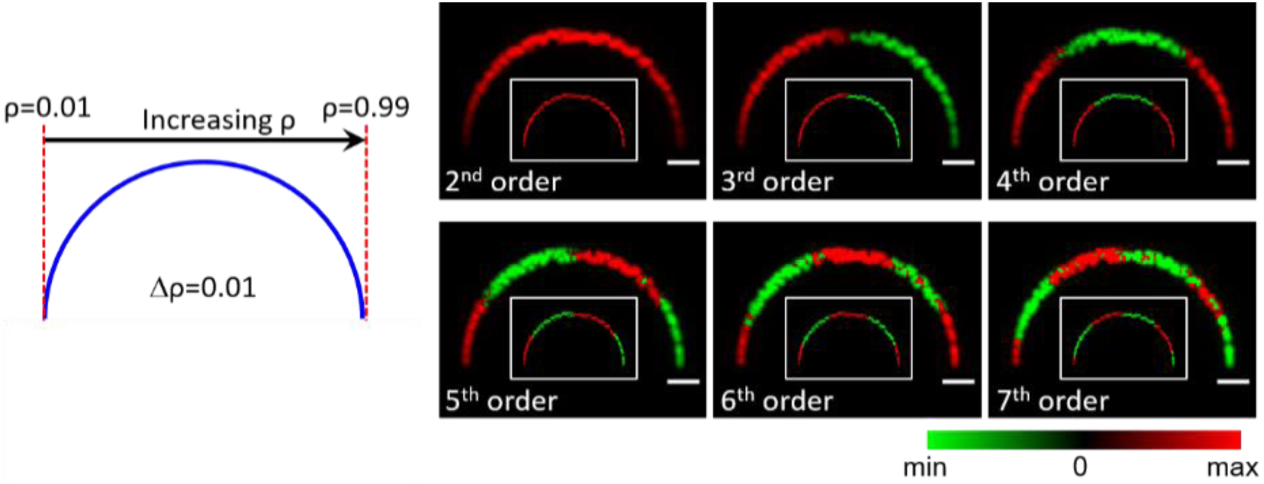
Simulations (‘Simulation-3’) to elucidate dependence of cusp artifacts on slowly varying ρ. A semicircle is populated with emitters having ρ values ranging from 0.01 to 0.99, as indicated in left panel, with interval being 0.01. Cumulants of different orders are displayed using gamma scale (gamma = 1/n) in right panels as labeled. Color coding is represented by the bar at bottom right (green for negative values, red for positive values, black for 0). In each image panel, inset shows ground-truth virtual emitters of given order, with red indicating positive virtual emitters and green indicating negative virtual emitters. Number of zero crossings (green/red transitions) increased with order of cumulants along gradient of ρ values, which is in agreement with ground truths shown in insets. Scale bars: 1.4 μm. Image intensities are displayed with gamma correction, with gamma value equals to the multiplicative inverse of the cumulant order.

To summarize this part: mixed populations of positive and negative virtual brightnesses cause cusp artifacts. Knowing the distribution of the virtual brightnesses based on the blinking statistics can allow one to predict cusp artifacts. SOFI cumulants that intrinsically have cusp artifacts are more vulnerable to background noise because of the lowering of the amplitude of the virtual signal. Such signal attenuation is owing to the neutralization of the positive and negative virtual brightnesses as well as the attenuation caused by the small amplitudes of the cumulant wn. Bleaching affects the ability to predict cusp artifacts; however, performing bleaching correction can help overcome this problem, making cusp artifact predictions valid again. Too small of an fbc value can make the SOFI cumulants more vulnerable to background noise (especially when the cumulant order is high) but does not have a significant effect on the statistical significance of the blinking trajectories. Lastly, and more interestingly, in the ideal cases, where the spatial variation of the blinking statistics is very small, cusps can serve as boundaries to segments with similar blinking statistics within the segment, but different blinking statistics between different segments.

## 6. Effects of cusp artifacts on image deconvolution and dynamic-range expansion correction and moment alternative

We have shown in the previous sections that spatial variations in the “on”-time ratio, □, lead to positive/negative-value variations in the cumulants, which, in turn, produce cusp artifacts once the absolute value operator has been used with the calculated cumulant images. The subsequent steps in the SOFI protocol for obtaining the final SOFI image include deconvolution (and/or Fourier reweighting) [18] and dynamic-range expansion correction (bSOFI[19], or ldrc as introduced in the accompanying manuscript [21]). In this section, we examine the effects of cusp artifacts on these two final (“post-processing”) steps. We argue that if cusp artifacts are not taken care of properly, their adverse effects could be amplified by the post-processing steps.

In order to determine the factors that contribute to the imperfections in deconvolution and Fourier reweighting, simulations were performed for an ideal collection of positive and negative virtual emitters (Fig. 7i) corresponding to a 3^rd^-order cumulant with zero background and noise; the results are shown in Fig. 7. The blurred image (Fig. 7ii) was generated by convolving the ground-truth image with the PSF. In the amplitude display (absolute value) of this image (Fig. 7iii), cusps can be seen clearly between the positive/negative regions. The images were generated on fine grids without noise, eliminating the imperfections which could arise from noise or pixilation, allowing us to analyze how the deconvolution step could be influenced by the cusp artifacts and the nature of mixed positive/negative virtual brightnesses. Notice that the imperfections caused by the discrete Fourier transform are still present in the image. As can be seen in the ideal deconvolution panel (Fig. 7iv), where under this ideal case, a perfect deconvolution is performed by direct division by the optical transfer function (OTF) in the Fourier space and the subsequent use of the inverse Fourier transform. In this case, we can see that even with a perfect imaging condition, we still expect minor imperfections (shown as the ringing artifacts in Fig. 7iv) caused by the discrete Fourier transform with finite boundaries. However, the virtual emitters are recovered in Fig. 7iv, with their virtual brightnesses remaining unchanged (including the negative ones), suggesting that the influence of discrete Fourier transform is negligible. In the case of Fourier reweighting (mimicking the case of 3^rd^-order cumulants), the transformed image was divided by a modified OTF with a damping factor (we used the machine epsilon variable in MATLAB, which represents a small number). The Fourier spectrum was subsequently multiplied by an OTF with a wider support. This was equivalent to convolving the ground truth with a smaller Gaussian PSF (Fig. 7v). Since the signs of the virtual brightnesses were conserved, the cusp artifacts remained in the high-density regions. On the other hand, when we performed deconvolution with solvers that imposed positivity constraints (for example, MATLAB’s “*deconvlucy*” or “*deconvblind*” functions), deconvolution failed (Fig. 7vi). Such failure is due to the presence of negative virtual brightnesses and cusps, which conflict with the positivity constraints used in the deconvolution solver. When the ground-truth is a mixture of positive/negative signal sources, the positivity constraint forces the pixel values to be positive either by taking only the absolute values or by rejecting all the negative ones during the recovery iterations, causing the image no longer to be the result of a convolution between a PSF and a ground-truth image, therefore deconvolution becomes fundamentally invalid.

**Fig. 7.**
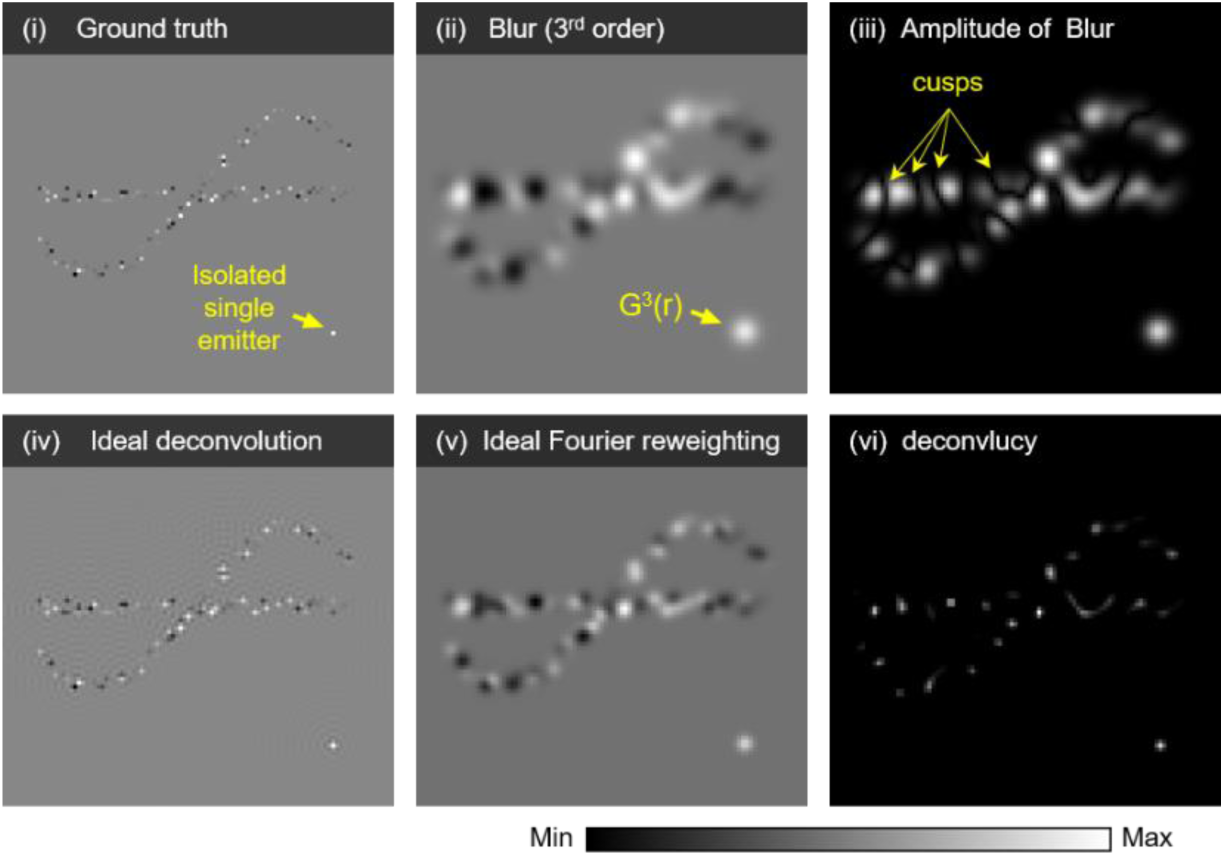
Post-processing of SOFI reconstructions containing cusp artifacts (simulation). Amplitudes of reconstructions are shown in grayscale (each panel has different dynamic range). Background of each panel is always zero (and therefore should be used as reference). Negative pixel values have darker colors than background; positive pixel values have lighter colors than background. (i) Ground-truth virtual emitters with both positive and negative values. (ii) Corresponding 3^rd^-order cumulant image (convolved with PSF). (iii) Amplitude (absolute value) of (ii) cusps are clearly visible. (iv) Ideal deconvolution result obtained by dividing Fourier-transformed image by optical transfer function (OTF) and subsequently performing inverse Fourier transformation. (v) Ideal Fourier reweighting, where, in contrast to the case for ideal deconvolution, Fourier spectrum is multiplied by extending the OTF before followed with inverse Fourier transform. (vi) Deconvolution result obtained using “deconvlucy” function, which imposes positivity constraint, that could affect the deconvolution when the corresponding ground-truth contains a mixture of positive and negative virtual brightnesses. PSF is simulated as perfect Gaussian with standard deviation of 4 pixels, as shown by the isolated emitter at the bottom right corner on each panel.

In cases where a subsequent dynamic range compression step (i.e., the “balancing” step in bSOFI) is performed after deconvolution, the final SOFI image will be even more distorted. Note that the current open source bSOFI package (MATLAB version [27]) uses “*deconvlucy*,” which imposes positivity constraints and therefore improperly handles negative virtual brightnesses. The reliance of bSOFI on positivity constraint and the assumption of perfect deconvolution should be removed. In contrast, Fourier reweighting does not alter the sign of the virtual brightnesses but is very sensitive to noise and does not eliminate cusps.

As shown in Eq (2.6) and discussed in Section 2, cumulants are additive. Therefore, the *n*^th^-order cumulant can be expressed as a sum of the cumulants of the individual emitters. This property is, in fact, the reason why cumulant reconstruction was originally chosen [9]. Reconstruction by moments (rather than by cumulants) was not considered owing to the presence of mixed terms that contain signal contributions from multiple individual emitters; such a reconstruction would be mathematically non-rigorous and the resulting physical meaning uninterpretable. Nonetheless, since even-order moments are intrinsically free of cusp artifacts, we pursued moments-based reconstruction as a ***practical*** approach and have shown that moments do result in an enhancement in the resolution as compared to the diffraction limit (as shown in the accompanying manuscript [21] and Section 8).

## 7. Comparison of performances

As shown in Fig. 8, we next compared the performances of SOFI, bSOFI, and 6^th^-order moments (M6) + ldrc for 6^th^-order reconstructions in terms of cusp artifacts.

**Fig. 8.**
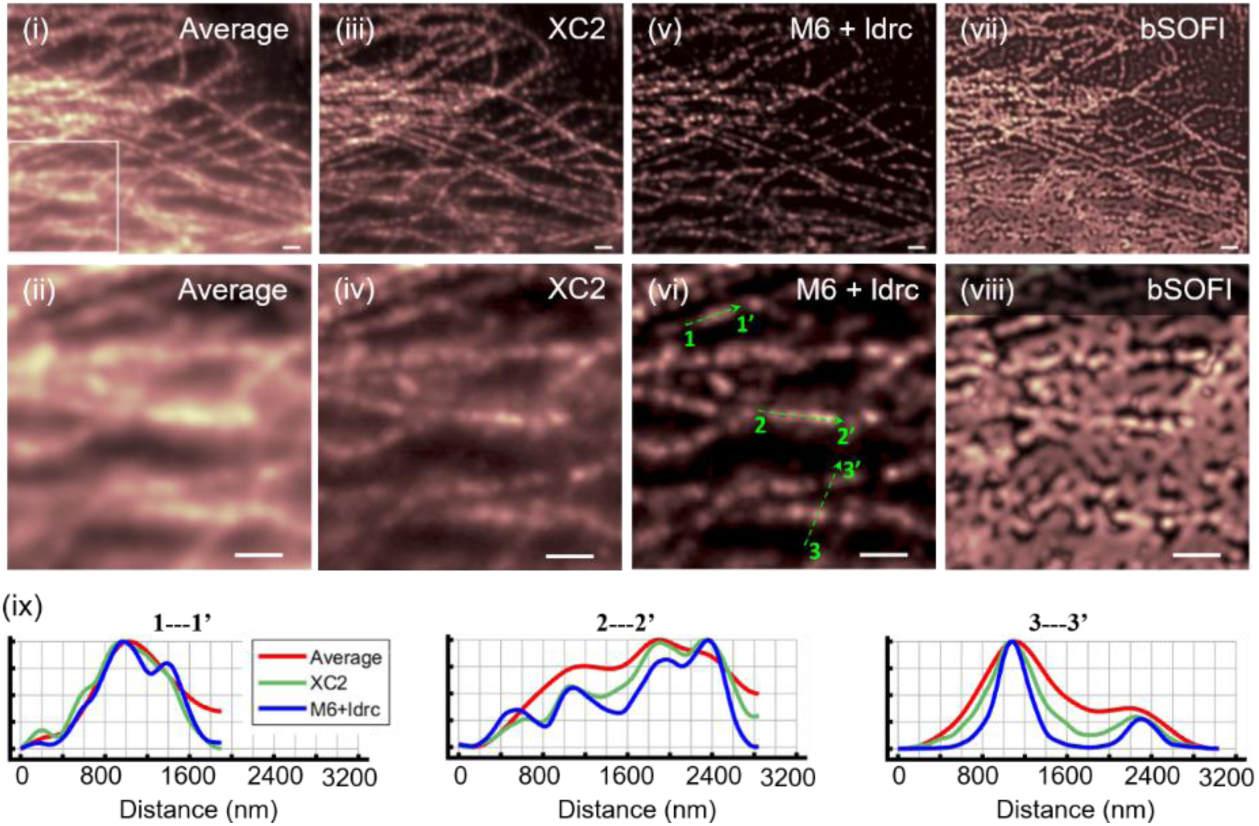
QD800-labeled microtubules (experimental data). For top two rows - <u>first column:</u> (i) average image of movie (2000 frames, 30 ms per frame) of QD800-labeled α-tubulin in fixed HeLa cell. (ii) magnified boxed region in sum image (i); <u>second column:</u> (iii) and (iv) show 2^nd^-order SOFI cumulants with extra pixels generated by cross-correlations (XC2), corresponding to panels (i) and (ii) respectively; <u>third column:</u> 6^th^-order moment-reconstruction with local dynamic range correction, corresponding to panels (i) and (ii) respectively; <u>fourth column:</u> 6^th^-order bSOFI corresponding to panels (i) and (ii). (ix) in last row: normalized intensity profiles of three green dashed lines (1---1’, 2---2’, and 3---3’) as labeled on top of each panel. Intensity profiles of Average, XC2, and M6+ldrc are compared; legend is provided in left-most panel. Detailed analysis of moments reconstruction are available in the accompanying manuscript [21].

All reconstructions were performed on the same data set (movie) of an α-tubulin network in a fixed HeLa cell that was labeled with blinking QD800 QDs (ThermoFisher Scientific, Carlsbad, CA). A total of 2000 movie frames were collected at 33 Hz using an EMCCD camera (Andor USA, South Windsor, CT). The results are shown in Fig. 8. Note that neither deconvolution nor Fourier reweighting was performed for M6 in order to isolate the factors responsible for the enhancement in the resolution and to assess the degree to which cusp artifacts degrade the overall performance. Average, XC2, M6 + ldrc, and bSOFI all show faithful reconstructions in the regions where the filament density is low and the α-tubulins are well separated. On the other hand, XC2 and bSOFI perform better than M6+ldrc in terms of feature visibility. However, in the regions where the filament density is high, as shown in the boxed region in (i) as well as in (ii), (iv), (vi), and (viii), M6-ldrc out-performs Average, XC2, and bSOFI. This is further confirmed by a comparison of the cross-sectional intensity profiles in (ix). In some cases, for M6+ldrc, two distinct peaks are observed, in contrast to the case for XC2 (see left-most panel in (ix)), suggesting that the degree of resolution enhancement was higher than that for XC2. However, we would still like to set the lower limit of resolution enhancement for M6+ldrc to be 2^1/2^, meaning that the resolution enhancement in this case should at least be similar to that for XC2. An additional theoretical *n*^1/2^-fold enhancement in the resolution for the *n*^th^-order moment can be realized when deconvolution is also used. A more detailed performance analysis of the moment-reconstruction method is presented in the accompanying manuscript [21]. Note that, for the bSOFI reconstruction, a deconvolution step was included because balanced cumulant reconstruction is a post-processing step performed after deconvolution.

## 8. Discussion and conclusions

Cusp artifacts had not been identified previously for several reasons: (i) 2^nd^-order cumulants are always positive and therefore are not susceptible to these artifacts; hence, many studies on SOFI stopped at the 2^nd^ order; (ii) post-processing steps such as balancing and deconvolution mask the existence of cusp artifacts and their origin; and (iii) SR methods in general, and SOFI and SOFI-derivatives reconstructions in particular, are an attempt to solve an ill-defined inverse problem (the location of the emitters in the sample). It is very hard, if not impossible, to identify cusp artifacts from experimental data. It was only after we had performed theoretical analysis of high order cumulants, realistic simulations (including those that considered dark noise, read-out noise, background noise, out-of-focus lighting, spatial variability in the photophysical properties with either limited or sufficient statistics, and the variation of the ground truth feature) and compared the results of the reconstruction algorithms to the ground truths were we able to successfully identify and characterize cusp artifacts. As shown in Fig. 3, once cusps had been identified, a careful reexamination of the experimental data did indeed confirm their existence.

Cusp artifacts are, in fact, hard to avoid. Even if the photophysical properties (blinking and photobleaching) of the emitters are more or less uniform across the sample, the finite acquisition time of a SOFI experiment (usually ~2,000 to 20,000 frames) is often not long enough to ensure that the statistical significance of the blinking behavior will be reached. Hence, the ρ values exhibit a broad distribution. This, in turn, leads to positive and negative higher-order (>2) cumulant values (Fig. 1). It is only when all the emitters in the sample exhibit a narrow ρ distribution during the data acquisition stage that cusp artifacts can, in principle, be avoided (as shown in Fig. 4 for P1 with cumulants of 2^nd^, 4^th^, and 6^th^ orders.). However, a narrow ρ distribution could still be positioned close to a zero crossing of one (or more) of the high-order cumulants, leading to the coexistence of both positive and negative cumulant values.

To minimize the adverse effects of cusp artifacts, the following guidelines should be considered: (1) emitters with uniform photophysical parameters (blinking and bleaching) should be used to ensure an intrinsically narrow ρ distribution; (2) long movies (with a large number of frames) should be acquired (to narrow down the experimental ρ distribution); (3) to the extent possible, the ρ distribution should be tuned to a zero-crossing-free zone (see Fig. 1); and (4) bleaching should be minimized and bleaching correction [14] should be performed on the data set, while following guidelines (1)-(3) for each bleaching correction block.

In summary, we successfully identified cusp artifacts in SOFI reconstructions with orders greater than two. These artifacts have either missed completely or interpreted erroneously in previous studies. In this work, we proposed a virtual-emitter interpretation of high-order SOFI cumulants, using which we could determine the theoretical origins of cusp artifacts. We performed a series of realistic simulations that provided insights into the origin of cusp artifacts and have proposed guidelines on how to minimize their adverse effects. We were able to demonstrate that moment-based reconstructions can improve the resolution and sometimes even eliminate these artifacts. We also suggest guidelines on how to screen for improved probes that could minimize cusp artifacts in high-order SOFI cumulants. In addition, based on the novel theoretical framework of virtual-emitter interpretation, our work provides insights into high-order cumulants as well as cusp artifacts that could potentially inspire positive utilization of the abnormal virtual brightness distribution and cusps where ρ can serve as an indicator.

## Author Contributions

S. W. and X. Y. designed the study; X. Y. performed the experiments, performed the simulations and analyses and wrote the associated codes; and S. W. and X. Y. wrote the manuscript. The authors declare no conflict of interests.

## Acknowledgements

The first author thanks Dr. Jianmin Xu for advises and training on cell culturing and immunostaining. We also thank Ms. Yingyi Lin, Mr. Xi Lin, and Mr. Sungho Son for their help with the project as undergraduate student researchers. Finally, we acknowledge the computational and storage services provided by the Hoffman2 Shared Cluster maintained by the Research Technology Group of the UCLA Institute for Digital Research and Education. This work was partially performed under the auspices of the U.S. Department of Energy by Lawrence Livermore National Laboratory under Contract DE-AC52-07NA27344.

