## Appendix for "Cusp-artifacts in high order Superresolution optical fluctuation imaging"

This document provides the appendix to "Cusp-artifacts in high order Super-resolution Optical Fluctuation Imaging". It contains more details for the derivations, simulation descriptions and comparison cases.

### 1. Appendix 1: Cumulants of single emitter blinking trajectory

Here we provide detailed derivations of Equation 4.1 in the main manuscript. We assume an emitter that stochastically switches between an 'on' (bright) state to an 'off' (dark) state with a time intensity trajectory F(t) = ϵb(t), where ϵ is the brightness of the emitter when it is in the 'on' state, $b(t)$ is the stochastic fluctuation profile, with $b(t)_{on} = 1$ and $b(t)_{off} = 0$. Given a finite time measurement, the fraction of time that the emitter stays in the 'on' state is given by ρ. We therefore have:

$$\langle b \rangle_t = \rho \, . \tag{SI-1.1}$$

The center-shifted fluctuations profile $b(t)$ is given by:

$$s(t) = \delta b(t) = b(t) - \langle b \rangle_t = b(t) - \rho \, . \tag{SI-1.2}$$

When the emitter is in the 'on' state, the center-shifted 'on' signal $s_{on}$ is given by:

$$s_{on} = b(t)_{on} - \langle b \rangle_t = 1 - \rho \, . \tag{SI-1.3}$$

Similarly, the center-shifted 'off' signal $s_{off}$ is given by:

$$s_{off} = b(t) - \langle b \rangle_t = -\rho \, . \tag{SI-1.4}$$

The n^th^ order moment ($m_n$) of $\delta b(t)$ will thus be:

$$\begin{aligned} m_n &= s_{on}^n \cdot \rho + s_{off}^n \cdot (1-\rho) \\ &= (1-\rho)^n \cdot \rho + (-\rho)^n \cdot (1-\rho) \end{aligned} \tag{SI-1.5}$$

The transformations between cumulants ($\omega_n$) and moments ($m_n$) (about the mean) are given by [23]:

$$
\begin{aligned}
\omega_2 &= m_2 \\
\omega_3 &= m_3 \\
\omega_4 &= m_4 - 3m_2^2 \\
\omega_5 &= m_5 - 10m_3m_2 \\
\omega_6 &= m_6 - 15m_4m_2 - 10m_3^2 + 30m_2^3 \\
\omega_7 &= m_7 - 21m_5m_2 - 35m_4m_3 + 210m_3m_2^2
\end{aligned}
\tag{SI-1.6}
$$

By substituting the expression of $m_n$ as a function of ρ from equation (SI-1.5) into equation (SI-1.6), and simplify the equations, we get:

$$
\begin{aligned}
\omega_2(\rho) &= \rho - \rho^2 \\
\omega_3(\rho) &= \rho - 3\rho^2 + 2\rho^3 \\
\omega_4(\rho) &= \rho - 7\rho^2 + 12\rho^3 - 6\rho^4 \\
\omega_5(\rho) &= \rho - 15\rho^2 + 50\rho^3 - 60\rho^4 + 24\rho^5 \\
\omega_6(\rho) &= \rho - 31\rho^2 + 180\rho^3 - 390\rho^4 + 360\rho^5 - 120\rho^6 \\
\omega_7(\rho) &= \rho - 63\rho^2 + 602\rho^3 - 2100\rho^4 + 3360\rho^5 - 2520\rho^6 + 720\rho^7
\end{aligned}
\tag{SI-1.7}
$$

which is the Equation 4.1 in the main manuscript.

### 2. Appendix 2: Cusp artifacts dependence on number of acquired frames

Here we discuss the dependence of cusp artifacts on total number of acquired frames (statistical significance). The 20k frames of P2 simulation-1 (see main text) were truncated to 10k, 4k, and 2k frames and SOFI cumulants were calculated respectively. Results are shown in Fig. S1. Predicted virtual emitters' signs are shown at the top of each column with (+/−) symbols. Cumulants from 2$^{nd}$ order to 7$^{th}$ order (using auto-correlations only) are plotted from left to right. Each row corresponds to the cumulants calculated from the given total number of frames indicated in the figure. For cumulant order where the virtual brightness exhibits a mixture of positive and negative values (6$^{th}$ order), no matter how many frames were processed, cusp-artifacts are always noticeable. This supports the notion that cusp artifacts are intrinsic to high order SOFI when the signs of the virtual brightness are mixed. For orders where virtual brightnesses have a pure sign (either positive or negative), as the total number of frames decreases, cusp artifacts start to show up. Also, the higher the cumulant order, the more frames are needed to reach statistical significance. In addition, at regions where there is out of focus light (shown with an arrow in Fig. S1(i)), cusp artifacts are enhanced.

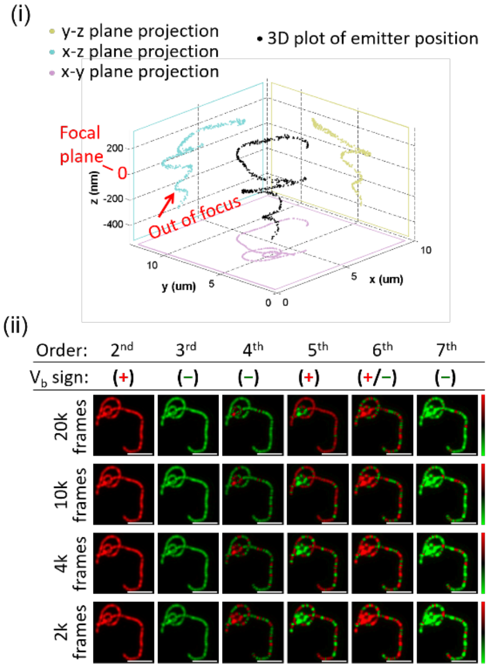


Fig. S1. Cusp artifacts dependence on number of acquired frames. (i) The ground-truth of emitters' locations in the simulation. The focal plane is at z = 0. A sub-population of emitters are places at > 200 nm out of focal plane to simulate the contribution of out-of-focus light. The Gibson Lanni's model was used to simulate the PSF with the following parameters: NA=1.4, wavelength = 800 nm, pixel size = 93.33 nm (detector pixel size 14 µm with 150x magnification). For these parameters, the axial PSF FWHM is 816.3 nm, while the PSF FWHM in x-y plane is 348.6 nm. (ii) shows the SOFI cumulants result with different total frames processed, at different cumulant order. All figures are gamma-corrected with $\gamma = 1/n$, where n is the cumulant order number. The predictions of the signs of the virtual brightnesses ($V_b$) are shown at the top. Scale bars: 2.8 um. Image intensities are displayed with gamma correction, with gamma value equals to the multiplicative inverse of the cumulant order.

### 3. Appendix 3: Cusp artifacts dependence on photobleaching and noise

Because the cusp artifacts and the associated cusp-artifacts predictions are affected by signal to noise ratio, and photobleaching (and corrections of photo-bleaching). Here in this section, we will first introduce the way we implemented the bleaching correction (3.1), and the way we simulated different signal levels of simulation data (3.2). The behavior of cusp-artifacts is then analyzed and compared under different photo-bleaching correction conditions and signal to noise level conditions in section 3.3 in this note.

#### *3.1 Implementation of bleaching correction*

Bleaching correction was implemented by truncating the overall movie series into individual blocks[14] with minor modifications. Here we provide detailed specifics about our implementation of bleaching correction used in this work with non-uniform block sizes for the movie truncation. Given a data set of N frames as shown in Fig. S2(i), the spatial average signal of each individual frame is calculated to derive a time evolution of the average signal $S_{ave}(t)$ as shown in Fig. S2(ii), then the high frequency components are filtered-out from $S^*_{ave}(t)$ to obtain a smoothed time evolution of average signal $S^*_{ave}(t)$ (Fig. S2(iii)). The overall averaged signal decrease of $S^*_{ave}(t)$ is calculated, and the movie is split into individual blocks where within each block, the amount of signal decrease is identical. This amount of signal decrease is characterized by the bleaching correction factor $f_{bc}$, which is the fractional signal decrease within each block (as compared to the total signal decrease over the whole movie). In Fig.

S2(iv), $f_{bc}$ was chosen to be $f_{bc} = 4\%$. In this example, the total movie is divided into 25 blocks, and within each block the signal decrease is 4% of the total signal decrease. Cumulants within each block are calculated independently, and averaged to obtain the final SOFI cumulants.

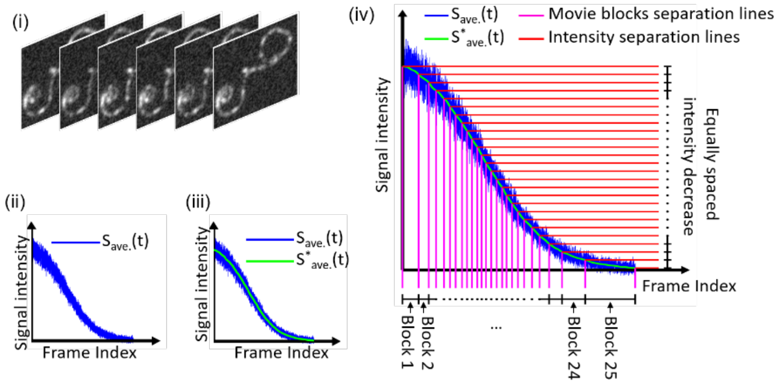


Fig. S2. Bleaching correction. Given a dataset consisting of multiple consecutive frames (i), the spatially averaged signal of each individual frame is calculated and the overall averaged time evolution $S_{ave}(t)$ is extracted, as in (ii). (iii) shows the smoothed time evolution of average signal $S^{*}_{ave}(t)$ after low-pass filtering. The entire movie is then split into individual blocks as shown in (iv). The total signal decrease within each block is imposed to be identical for all blocks (as determined from the smoothed time evolution $S^{*}_{ave}(t)$ and the bleaching correction factor $f_{bc}$). Panel (iv) demonstrates how the movie was split into 25 blocks (of different sizes.) with $f_{bc}$=0.4%.

#### *3.2 Cusp artifacts dependence on different signal-to-noise (S/N) ratio*

Different signal to noise ratios are simulated to study the dependence of cusp artifacts. We first simulated data without background noise. The signal itself, governed by Poisson distribution, was simulated by resetting the signal value with a random number generated from Poisson distribution where the expectation value was the true signal value, denoted here as $s(t)$. Real (experimental) background noise (denoted here as $n(t)$) was then added to $s(t)$. For this, we recorded frames under bright (uniform) illumination conditions (Fig. S3) with an empty sample. The simulated movie $s(t)$ was multiplied by different constant values $c$ to simulate different signal level $c{\cdot}s(t)$. The background noise $n(t)$ was then added to result in a total signal of $c{\cdot}s(t)+ n(t)$. Fig. S3 shows the strategy of simulating two different signal (and S/N) levels: $6{\cdot}s(t)+ n(t)$ in (ii) and $3{\cdot}s(t)+ n(t)$ in (iii). Histograms of signal ($M$) pixels values versus background ($M'$) pixels values are then plotted. The corresponding calculated peak signal to noise ratio (SNR) are 1.47 and -1.33 as calculated according to the formula below:

$$SNR_{dB} = 10 \cdot \log_{10}\left(\frac{P_{signal}}{P_{noise}}\right) \qquad \text{(SI-1.8)}$$

Where $P_{noise}$ is the average power of the background noise, $P_{signal}$ is the average power of the signal. As shown in Fig. S3, the masked region contains both signal and noise component, while the region outside of the mask contains only the noise component. Therefore, $P_{noise}$ is calculated as the average pixel value outside the masked region, and $P_{signal}$ is calculated as the average pixel value inside the masked region followed by subtraction of $P_{noise}$.

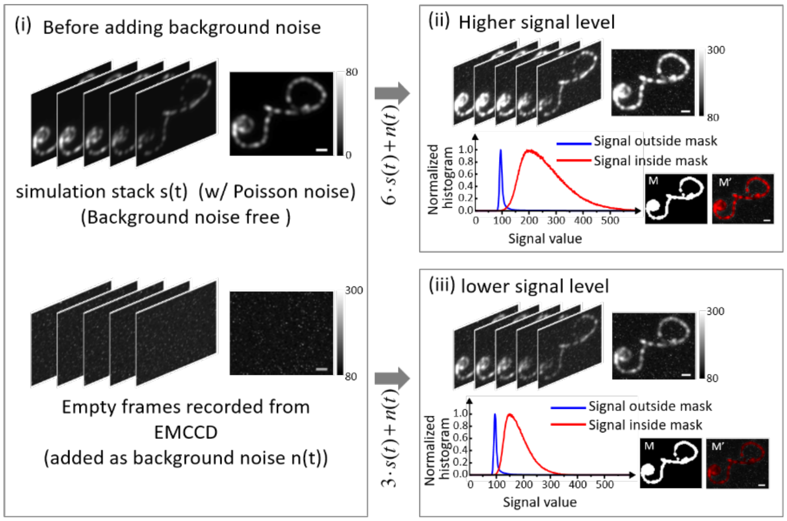


Fig. S3. Simulation with different S/N. (i) shows two movie stacks, the top movie stack $s(t)$ represents the simulated noise-free movie (of blinking probes). The recorded background $n(t)$ is added to $s(t)$ as the background noise. (ii) shows the constructed stack of noisy movie using $6s(t)+n(t)$; (iii) shows constructed stack of noisy movie using $3s(t)+n(t)$. Pixel intensity histograms for (ii) and (iii) are shown on the right panels. For each panel, *M* represents a mask that blocks the simulated morphology and therefore used for calculating the background signal histogram (blue) and *M'* represents a mask that blocks all background pixels (outside simulated morphology) and therefore used for calculating the signal histogram (red). Histograms were calculated from pixels of 1000 simulated frames. Scale bars: 840 nm.

#### *3.3 Cusp artifacts dependence on photobleaching and noise*

Effects of bleaching were tested first for the dataset without background noise (Fig. S4), for bleaching correction factors $f_{bc}$=0.25%, 0.5%, 1%, 4%. Photobleaching simulations demonstrate that bleaching can alter the virtual brightness distribution (to deviate from predictions based on blinking statistics). Bleaching correction, however, can recover the virtual brightness distribution to the case without bleaching that agrees with the prediction based on blinking statistics when sufficient statistical significance is provided. Note here that bleaching correction can recover the theoretical virtual brightness distribution, but it is not designed to directly reduce cusp artifacts. To further examine how noise can affect the cusp-artifacts predictions, two different simulations with two different signal levels were generated with the scheme depicted in Fig. S3. SOFI cumulant calculations were performed with the same choices of $f_{bc}$ as shown in Fig. S4. Results are displayed in Fig. S5. SOFI cumulants that exhibit cusp artifacts are more vulnerable to the addition of background noise. This is because in cases where cumulants exists, the signal amplitude is also attenuated, therefore more vulnerable to the background noise.

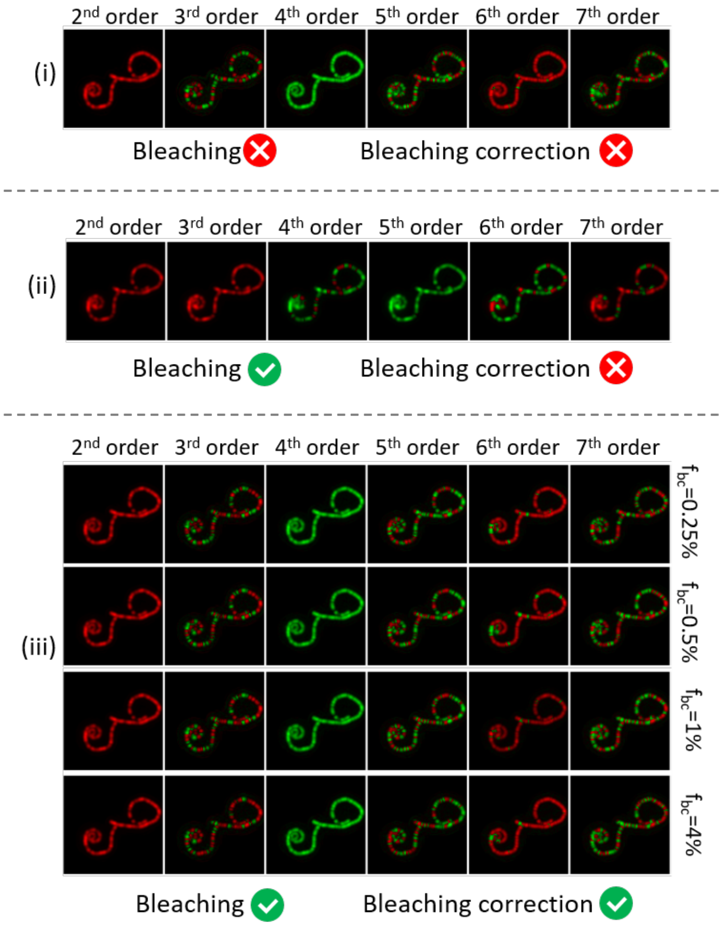


Fig. S4. Effect of bleaching correction and different choices of factors on cusp artifact. Each panel in this figure shows the Gamma-corrected SOFI cumulants from 2nd order to 7th order are plotted from left to right display with gamma correction factor of $\gamma = 1/n$, where n is the cumulant order. (i) shows the result from the P1 population in 'simulation-1'. In this simulation there is no bleaching effect therefore no bleaching correction is needed. We can see that the virtual brightness sign follows the predicted distribution shown in Figure 2. (ii) shows the SOFI processed results of 'simulation-2', which is generated by adding bleaching effect to the P1 population in 'simulation-1'. Bleaching correction is not performed and we can see that the virtual brightness deviates from the theoretical predictions, and the cusp-artifacts behaves different than that in panel (i). We then performed bleaching correction before the SOFI processing with different bleaching correction factors: $f_{bc}$ = 0.25%, 0.5%, 1%, 4% (panel (iii)). With bleaching correction, the virtual brightness and the feature of the cusp-artifacts are restored and becomes similar with the case without bleaching as shown in panel (i). And such restoration is not sensitive to the choice of this bleaching correction factor. Image intensities are displayed with gamma correction, with gamma value equals to the multiplicative inverse of the cumulant order.

Interestingly, the 4th order cumulants exhibit pure negative virtual brightness values. Under such conditions, the feature of interest is more visible and exhibit enhanced contrast (negative contrast) against the background which follows Poisson distribution. This is because cumulants of the center-shifted Poisson random variable is always positive (see below).

### 4. Appendix 4: High order cumulants with Poisson background noise

Here we discuss the analytical form of high order SOFI cumulants (for auto-correlations without time lags) considering the captured movie being corrupted by a noise component that follows Poisson distribution.

High order cumulants (for orders greater than one) of a center-shifted Poisson random variable is always λ (shown in 4.1 below). Additionally, cumulants are additive [23]. Therefore, when a component of additive background noise is considered, the corresponding contribution to the SOFI cumulant can be separated from the major summation notion and serve as a positive and additive term (λ) to the analytical form of SOFI cumulants.

#### *4.1 Cumulant of Poisson random variable*

We note here that for SOFI analysis, the first step is to 'center-shift' each pixel value, defined as the subtraction of the average image (time-average) from every individual frame of the acquired movie. Specifically, we center-shifted the intensity time trace of each pixel independently, therefore the discussion of cumulants of the noise is also concerning the center-shifted noise component as a random variable.

Assume the noise σ follows Poisson distribution with constant expectation value λ:

$$P_\sigma(k) = \frac{\lambda^k e^{-\lambda}}{k!} \ , \qquad \text{(SI-1.9)}$$

where $k$ is a positive integer, and λ is the expectation value of σ (which is also a positive integer). $P_\sigma$ is the probability density function of σ. The center-shifted noise detection (denote as δσ) would be:

$$\delta\sigma = \sigma - \lambda \ , \qquad \text{(SI-1.10)}$$

The relationship between σ and δσ indicates that the probability of δσ= $k$ (expressed as $P_{\delta\sigma}(k)$) is equivalent to the probability of $\sigma$ =$k$+λ (expressed as $P_\sigma(k+\lambda)$). Therefore, the probability density function of δσ would be:

$$P_{\delta\sigma} = P_\sigma(k+\lambda) = \frac{\lambda^{(k+\lambda)} e^{-\lambda}}{(k+\lambda)!} \ . \qquad \text{(SI-1.11)}$$

and the moment-generating function of δσ (denoted as $M_{\delta\sigma}(t)$) can be deduced as:

$$\begin{aligned} M_{\delta\sigma}(t) &= E[e^{tk}] \\ &= \sum_{k=-\lambda}^{\infty} e^{tk} P_{\delta\sigma}(k) \\ &= \sum_{k=-\lambda}^{\infty} e^{tk} \cdot \frac{\lambda^{(k+\lambda)} e^{-\lambda}}{(k+\lambda)!} \\ &= e^{\lambda(e^t - t - 1)} \end{aligned} \ . \qquad \text{(SI-1.12)}$$

The cumulant-generating function (denoted as $K_{\delta\sigma}(t)$) is derived from the natural logarithm of $M_{\delta\sigma}(t)$:

$$K_{\delta\sigma}(t) = \ln M_{\delta\sigma}(t) = \lambda(e^t - t - 1) \ . \qquad \text{(SI-1.13)}$$

Take the Maclaurin series of the expression above, we get:

$$K_{\delta\sigma}(t) = 0 \cdot \frac{t^1}{1!} + \lambda \cdot \frac{t^2}{2!} + \lambda \cdot \frac{t^3}{3!} + \lambda \cdot \frac{t^4}{4!} \cdots \ , \qquad \text{(SI-1.14)}$$

where the front factor for the term of 1st power of $t$ is zero, and all the rest of the front factors are $\lambda$.

Because the n<sup>th</sup> power of $t$ is equivalent to the n<sup>th</sup> order cumulant, we conclude that for a random variable that follows Poisson distribution with mean value $\lambda$, the 1st order center-shifted cumulant equals to 0, all the higher orders of center-shifted cumulants are positive and equal to $\lambda$. Interested readers are suggested to M. G. Kendall's work [23] for more details concerning the properties of moments, cumulants as well as the corresponding generating functions.

### 5. Appendix 5: comparison of cumulants of order 2 to 7 with different image displays.

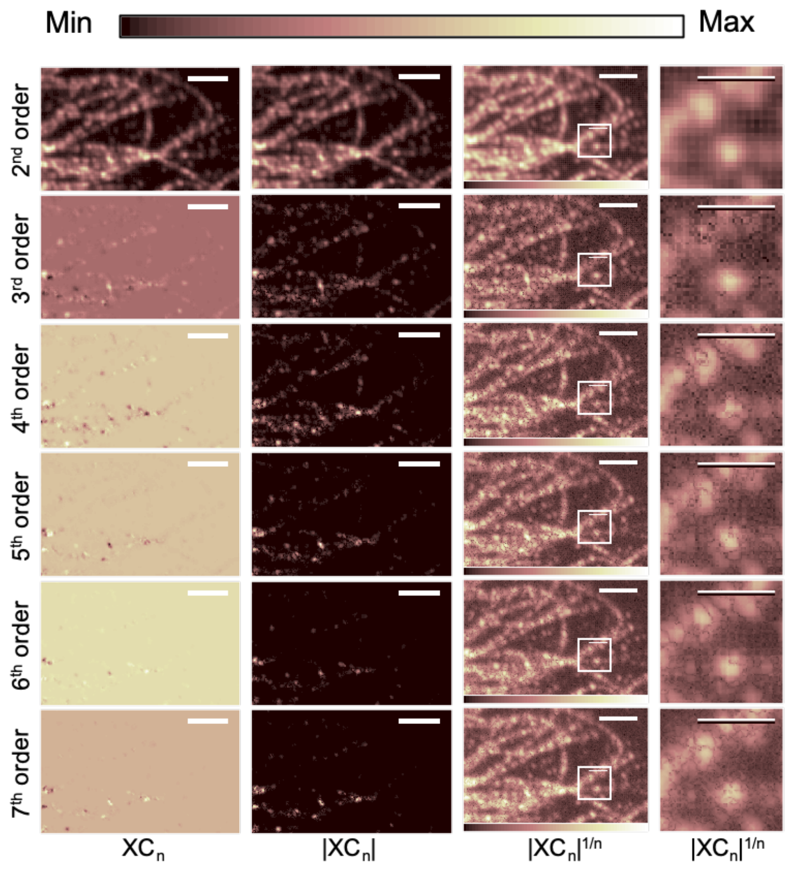


Fig. S5. we provide the cumulants from 2nd to 7th order with different image display schemes, including the original intensity range display (first column), absolute value of the original intensity display (second column), gamma corrected absolute value of the original intensity display (third column), a zoom-in panel of the third column shown with a box (fourth column). Scale bars: 3.2 μm for the first, second and third columns, and 1.6 μm for the fourth column.

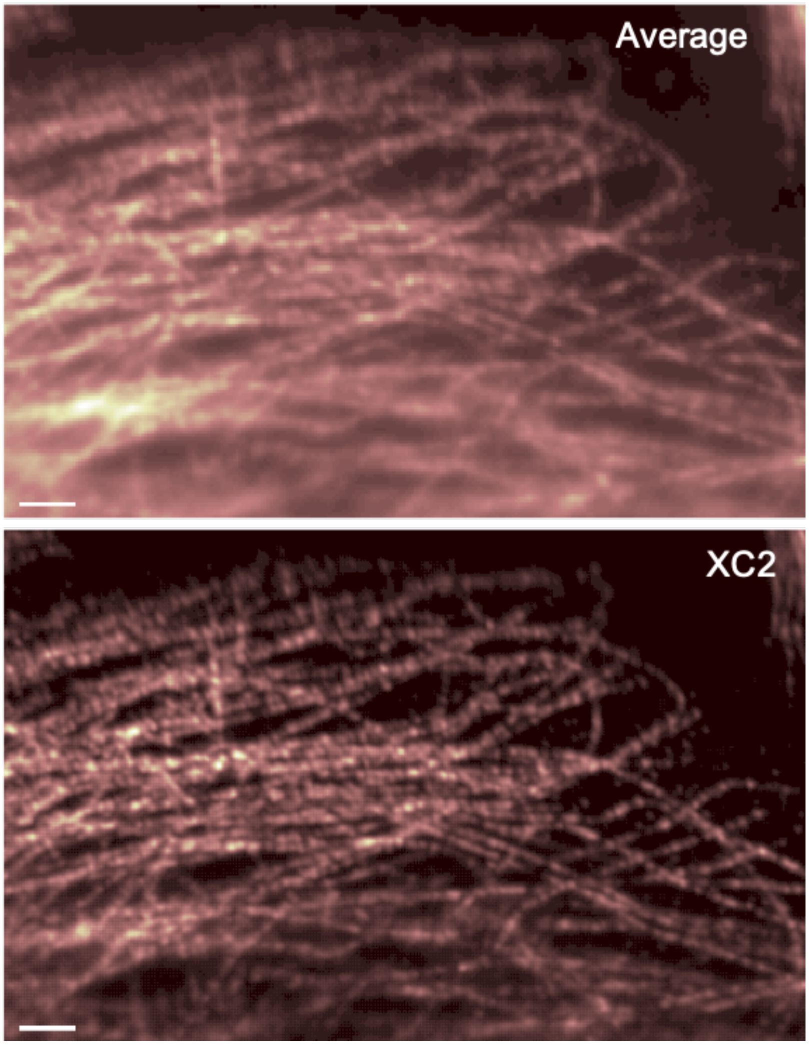


Fig. S6. a comparison of the average image to the second order cross-correlation SOFI cumulants to demonstrate the resolution enhancement of SOFI cumulant at second order. Scale bars: 3.2 μm.
